# Different functions of Lhcx isoforms in photoprotective mechanism in the marine diatom *Thalassiosira pseudonana*

**DOI:** 10.1101/2024.04.16.589823

**Authors:** Mana Nakayasu, Seiji Akimoto, Kohei Yoneda, Soichiro Ikuta, Ginga Shimakawa, Yusuke Matsuda

## Abstract

Photosynthesis needs light energy, but that exceeding the maximal capacity of photosynthesis enhances formation of reactive oxygen species, which potentially causes photodamages. Therefore, light-harvesting complexes (Lhc) in phototrophs harbor various proteins and pigments to function in both light capture and energy dissipation. Diatom Lhcx proteins are reported to be a critical component for thermal dissipation of excess light energy, but the molecular mechanism of photoprotection is still not fully understood and the functions of each Lhcx isoform are not yet differentiated. Here, we focused on two types of Lhcx isoforms in *Thalassiosira pseudonana*: TpLhcx1/2, putative major components for energy-dependent fluorescence quenching (qE); and TpLhcx6_1, functionally unknown isoform uniquely conserved in Thalassiosirales. TpLhcx1/2 proteins accumulated more under high light than under low light, while the TpLhcx6_1 protein level was constitutive irrespective of light intensities and CO2 concentrations. High-light induced photodamage of photosystem II was increased in the genome-editing transformants of these Lhcx isoforms relative to the wild-type. Transformants lacking TpLhcx1/2 showed significantly lowered qE capacities, strongly suggesting that these proteins are important for the fast thermal energy dissipation. While in contrast, genome-editing transformants lacking the TpLhcx6_1 protein rather increased the qE capacity. TpLhcx6_1 transformants were further evaluated by the low-temperature time-resolved chlorophyll fluorescence measurement, showing the longer fluorescence lifetime in transformants than that in the wild type cells even at the dark-acclimated state of these cells. These results suggest that TpLhcx6_1 functions in photoprotection through non-photochemical energy dissipation in the different way from qE.

**One sentence summary:** The marine diatom *Thalassiosira pseudonana* dissipates excess light energy for photoprotection *via* two types of mechanisms supported by different Lhc isofoms.

## Introduction

Diatom is a highly diverse group of algae with an estimate of >100,000 species (Mann and Vanormelingen 2013) and plays a major role for primary production at a global scale, resulting in 40% of total oceanic productivity which corresponds to 20% of carbon fixation on the earth (Levitan et al. 2014). Marine diatoms are widely distributed in oceanic environments, including the intertidal zone and the ocean surface, where the light intensity always fluctuates and the diatoms easily suffer from strong light exceeding the demand for photosynthetic CO2 fixation. Excess light energy has the potential to cause the generation of reactive oxygen species such as singlet oxygen and superoxide anion radical (Krieger-Liszkay and Shimakawa 2022). Therefore, diatoms develop the regulatory mechanism to dissipate the light energy as heat at the fucoxanthin-chlorophyll (Chl) *a*/*c* light-harvesting complex (FCP), which is the so-called qE observed as a non-photochemical quenching (NPQ) in Chl fluorescence analyses (Lepetit et al. 2012). There is a plausible consensus that diatom qE is supported by three factors: (1) de-epoxidation state of xanthophylls; (2) thylakoid lumen acidification; and (3) Lhcx proteins (Goss and Lepetit 2015). In one model, xanthophyll de-epoxidases are activated by thylakoid lumen acidification, and the de-epoxidized xanthophylls (diatoxanthin and zeaxanthin) dissipate the excitation energy originated from Chls as heat (Goss and Lepetit 2015). Lhcx proteins are suggested to support a process of energy transfer between Chls and xanthophylls in the pennate diatom *Phaeodactylum tricornutum* (Buck et al. 2019). However, either FCPs detached from photosystem II (PSII) core or bound to it are suggested to function for qE (Miloslavina et al. 2009; Chukhutsina et al. 2014; Lavaud and Goss 2014; Goss and Lepetit 2015; Giovagnetti and Ruban 2017). To date, several molecular models have been proposed for the non-photochemical fluorescence quenching in diatoms. Further, these mechanisms are likely to include species-dependent diversity. Overall, the molecular mechanism of diatom qE is still under debate.

Lhcx is one of fucoxanthin-Chl *a*/*c* binding (Lhc) protein groups that constitute the FCP family in stramenopiles and haptophytes. Based on phylogenetic analyses, diatom Lhc proteins are categorized into Lhcf, Lhcr, Lhcx, Lhcq, Lhcy, and Lhcz (Dittami et al. 2010; Nymark et al. 2013; Nagao et al. 2020; Kumazawa et al. 2022). Whereas Lhcf and Lhcr are major components of FCP to function as antenna proteins, Lhcx is grouped into LI818 clade like the stress-related green algal Lhc protein (Lhcsr). The mutant of the green alga *Chlamydomonas reinhardtii* lacking Lhcsr3 shows the lower qE capacity than the wild type (WT) that in turn resulted in the decrease in maximum efficiency of PSII photochemistry (Fv/Fm) in a high light condition, indicating that Lhcsr plays a role for photoprotection of PSII *via* qE (Allorent et al. 2013). In the diatom *P. tricornutum*, the lack of Lhcx impaired the qE capacity (Buck et al. 2019). These facts suggest that Lhcx is important for PSII photoprotection in diatoms. The marine centric diatom *Thalassiosira pseudonana* encodes six Lhcx genes: *TpLhcx1* (THAPSDRAFT_264921), *TpLhcx2* (THAPSDRAFT_38879), *TpLhcx4* (THAPSDRAFT_270228), *TpLhcx5*(THAPSDRAFT_31128), *TpLhcx6* (THAPSDRAFT_12097), and *TpLhcx6_1* (THAPSDRAFT_30385). From transcript analyses, TpLhcx1 seems to be the dominant Lhcx isoform in *T. pseudonana* (Zhu and Green 2010). Since TpLhcx2 shows an extremely high score of BLAST identity (>99%) to TpLhcx1, hereafter we term them altogether as TpLhcx1/2. TpLhcx1/2, TpLhcx4, and TpLhcx6 are highly expressed under high light (Zhu and Green 2010). Further, TpLhcx1/2 and TpLhcx5 have been suggested to interact with PSII together with associating diadinoxanthin de-epoxidase (Kansy et al. 2020). TpLhcx6_1 has been often detected in proteome analyses and suggested to interact with photosystem I (PSI), PSII, and FCP trimer (Grouneva et al. 2011; Calvaruso et al. 2020). In the centric diatom *Cyclotella meneghiniana*, the Lhcx6_1 ortholog is constitutively accumulated in FCP trimer (Gundermann et al. 2019). The latest work uncovered the structure of PSII-FCP supercomplex at 2.68-Å resolution by cryo-electron microscopy in *T. pseudonana*, demonstrating that TpLhcx6_1 binds to PSII (Feng et al. 2023). The structure suggested that TpLhcx6_1 itself binds 11 Chls and 3 carotenoids, similarly to other Lhc isoforms, and is located between PSII and the FCP dimer to optimize the excitation-energy transfer between them and also energy quenching at the site close to PSII (Feng et al. 2023). In spite of these biochemical and structural knowledges, the physiological functions of Lhcx6_1 isoforms have not yet been studied.

Here, we investigated the physiological functions of two types of Lhcx isoforms in *T. pseudonana* using the genome editing transformants of TpLhcx1/2 and TpLhcx6_1. Whereas the transformant phenotype clearly indicated that TpLhcx1/2 are main components for qE, the knock-out transformants of TpLhcx6_1 rather showed the larger qE capacity than WT, although both transformants suffered from PSII photodamage. In time-resolved Chl fluorescence measurements, we found that the fluorescence lifetime in the TpLhcx6_1 defective cells was longer in the transformants than in WT even in the dark-acclimated cells, suggesting that TpLhcx6_1 functions for photoprotection in the different manner from the generally-hypothesized mechanism of qE.

## Results

### Similarity among Lhcx isoforms in marine diatoms

Here, we rearranged the similarity among Lhcx isoforms in diatoms, including *T. pseudonana*, *P. tricornutum*, *Pseudo-nitzschia multistriata*, *Fistulifera solaris*, and *Minidiscus variabilis*, with the Lhcsr and PsbS isoforms in the green alga *C. reinhardtii*, the liverwort *Marchantia polymorpha*, the moss *Physcomitrella patens*, and the angiosperm *Arabidopsis thaliana*. PsbS is the Lhc-like PSII protein functioning for qE in land plants (Li et al. 2002). The phylogenetic tree mainly showed the three groups branched almost at the root (Fig. 1): (1) the clade containing Lhcx and Lhcsr isoforms in a variety of photosynthetic organisms; (2) the clade of the Lhcx6_1 isoforms; and (3) the clade composed of PsbS isoforms. Lhcx6_1 isoform was recognized only in the centric diatoms *T. pseudonana* and *M. variabilis*, while the other Lhcx isoforms were found in all the five diatom species. Although the coding sequence is not fully determined, Lhcx6_1 is also found in the genome of the centric diatom *Cyclotella cryptica* (https://phycocosm.jgi.doe.gov). Meanwhile, the centric diatom *Chaetoceros gracilis* does not possess Lhcx6_1 isoform (Kumazawa et al. 2022). The three diatom species *T. pseudonana*, *M. variabilis*, and *C. cryptica* are evolutionarily close around the order Thalassiosirales in diatoms (Arsenieff et al. 2020; Hevia-Orube et al. 2016).

**Figure 1.**
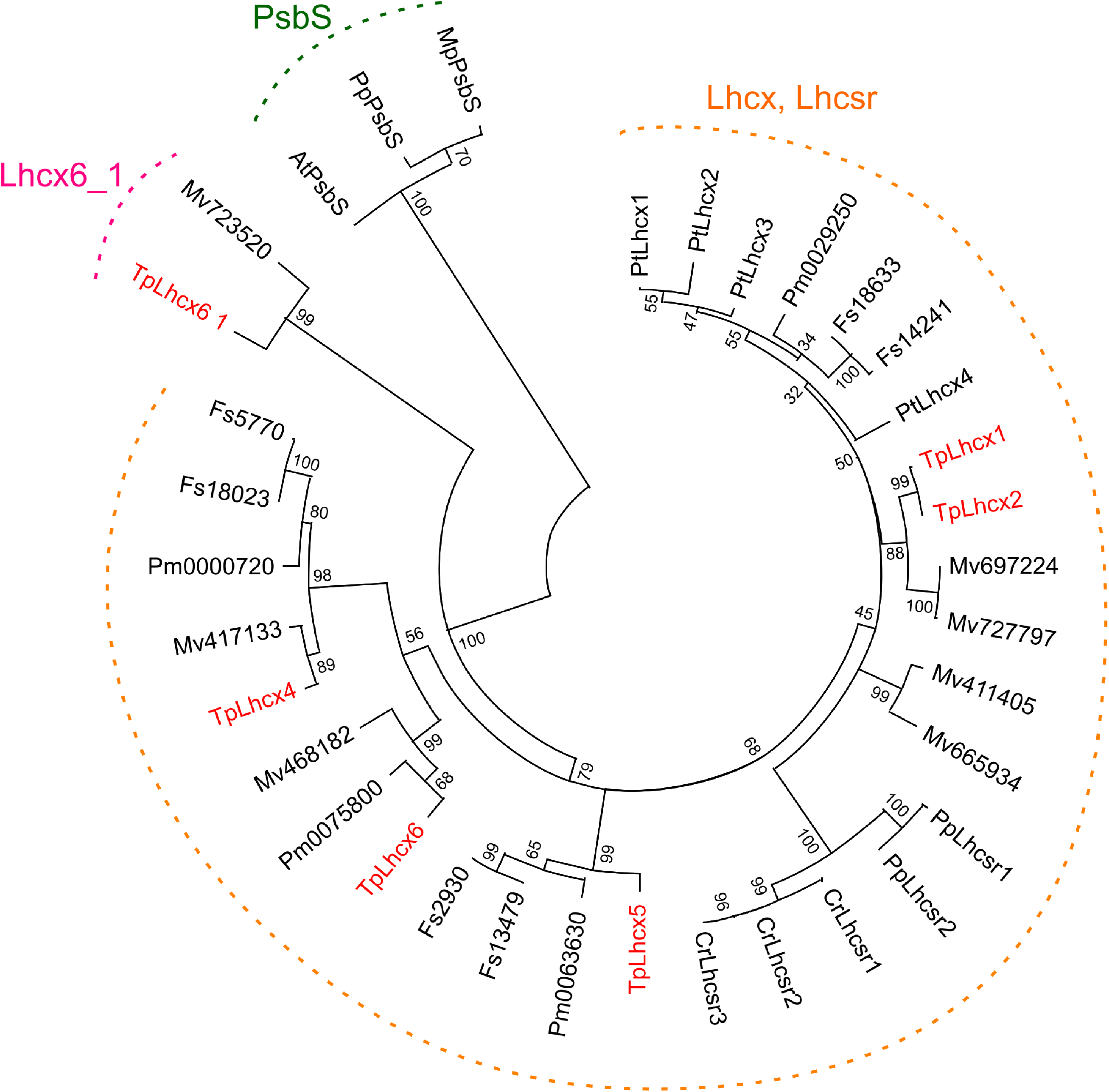
Maximum likelihood phylogenetic tree of Lhcx and its orthologs in photosynthetic organisms. A consensus tree was generated using 1000 bootstraps. The clade containing TpLhcx6_1 was distinguished from the other Lhcx and Lhcsr. The Lhc-like PSII protein (PsbS) functions as the critical component for qE in land plants. At, *Arabidopsis thaliana*; Cr, *Chlamydomonas reinhardtii*; Pp, *Physcomitrella patens*; Mp, *Marchantia polymorpha*; Pt, *Phaeodactylum tricornutum*; Pm, *Pseudo-nitzschia multistriata*; Fs, *Fistulifera solaris*; Tp, *Thalassiosira pseudonana*; Mv, *Minidiscus variabilis*. TpLhcx isoforms are highlighted in red. Accession numbers for each gene are listed in Supplemental Table S1.

Global distribution of TpLhcx1/2 and TpLhcx6_1 homologs was investigated using the meta-genomics/transcriptomics database (Villar et al. 2018). Both genes are broadly distributed in the oceanic environments, which should be related to the wide habitat of marine diatoms (Supplemental Fig. 1A). Whereas homologs to TpLhcx1 (E-value: < 10^−50^) were detected in a variety of micro algae, those to TpLhcx6_1 (E-value: < 10^−50^) was almost limited in the order Thalassiosirales (Supplemental Fig. 1B). Overall, Lhcx6_1 is the unique Lhcx isoform conserved in the diatom order Thalassiosirales.

### Accumulation of TpLhcx1/2 and TpLhcx6_1 proteins at different light and CO2 availability

Next, we compared the accumulation of TpLhcx1/2 and TpLhcx6_1 proteins in *T. pseudonana* among the cells grown under different light and CO2 conditions. Accumulation level of TpLhcx1/2 significantly increased in 2 h after the cells were shifted from low light (LL; 50 µmol photons m^−2^ s^−1^) to high light (HL; 500 µmol photons m^−2^ s^−1^) conditions (Fig. 2A and B), which was consistent to the recent report (Calvaruso et al. 2022). Meanwhile, TpLhcx6_1 showed little difference in the accumulation level between LL and HL conditions (Fig. 2A and C). It has been reported that CrLhcsr3 is highly expressed under low CO2 (LC) than high CO2 (HC) conditions (Ruiz-Sola et al. 2023). However, both TpLhcx1/2 and TpLhcx6_1 proteins showed no response in accumulation level by changing the growth CO2 condition (Fig. 2). These data indicated that the expression of the TpLhcx6_1 protein is constitutive regardless of light and CO2 availability, while the expression of TpLhcx1/2 is stimulated in HL conditions.

**Figure 2.**
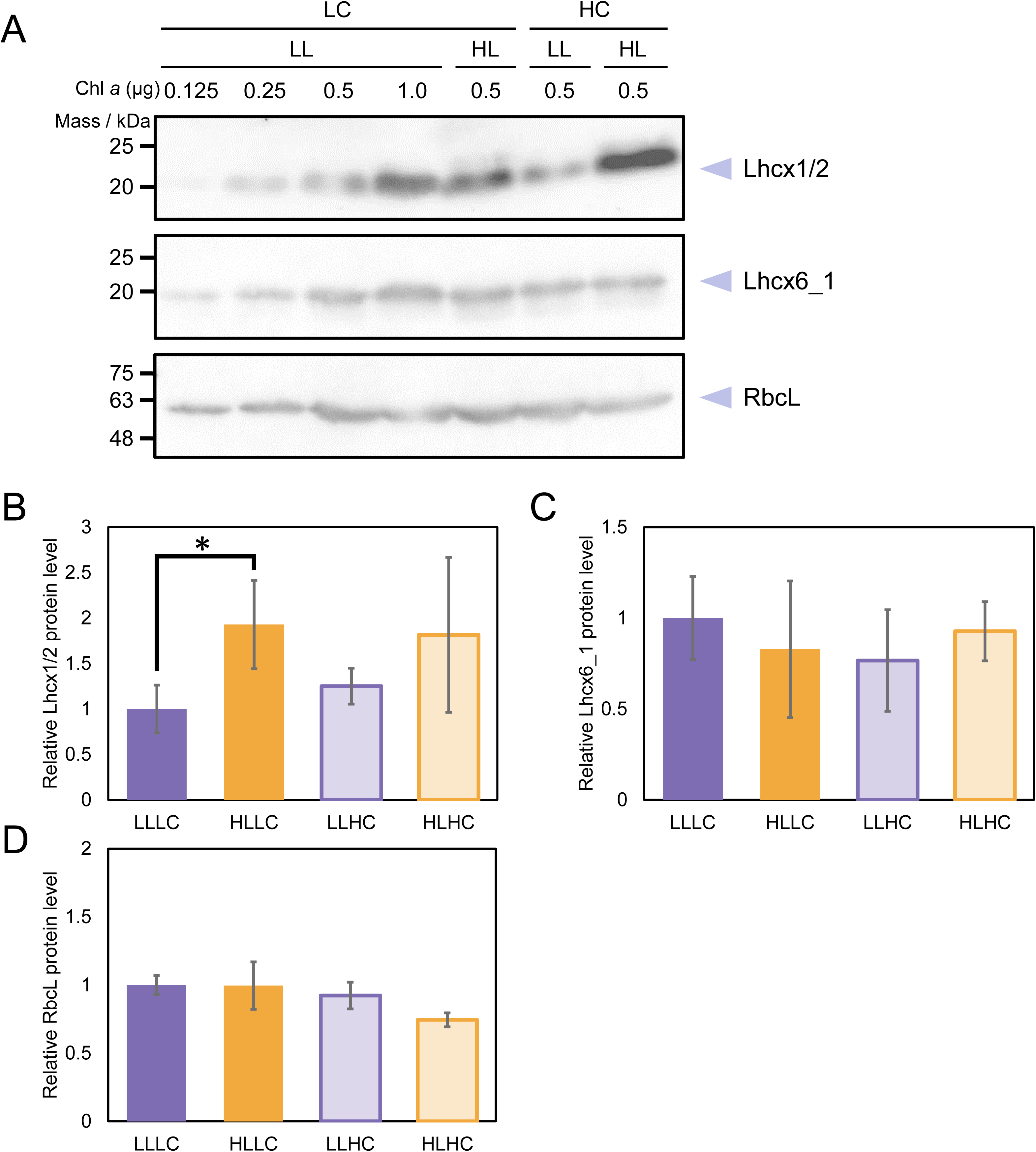
Levels of TpLhcx1/2 and TpLhcx6_1 protein detected by immunoblotting at different light intensity or CO2 concentration in *Thalassiosira pseudonana*. Whole crude extracts were prepared from the cells grown under air-level and 1% CO2 (LC and HC, respectively) before and after the 2-h high light (500 µmol photons m^−2^ s^−1^) exposure (LL and HL, respectively). (A) Representative blots are shown. RbcL was a loading control. (B−D) Levels of each protein was semi-quantified by densitometry. Data are shown in the mean with the standard deviation (*n* = 3, biological replicates). Asterisk shows the statistically significant difference (*p*<0.05) between LL and HL cells as per Student’s *t* test.

### Genome editing of TpLhcx1/2 and TpLhcx6_1

To assess the physiological functions of each Lhc isoform, we constructed the knock-out transformants of TpLhcx1/2 and TpLhcx6_1 in *T. pseudonana* by a highly specific genome editing, CRISPR-Cas9 nickase. For *TpLhcx1/2* genes, a pair of single guide RNAs (sgRNAs) were designed around the start codons (Fig. 3A and Supplemental Fig. S2). Due to the extremely high similarity (>99%), the sgRNAs to *TpLhcx1* also targets *TpLhcx2*. Finally, two independent transformant lines were established as the *TpLhcx1/2* double knock-out transformants (denoted as x1/2KO-1a and x1/2KO-1b). For *TpLhcx6_1* gene, two pairs of sgRNA were designed around the start codon and at the Chl binding domain (Fig. 3A and Supplemental Fig. S2). One each transformant harboring independent indels were obtained on two different target sites (denoted as x6_1KO-1 and x6_1KO-2). The lack of each Lhcx isoform was confirmed by immunoblotting with the specific antibodies to TpLhcx1/2 and TpLhcx6_1 (Fig. 3B) without any cross reaction between each Lhcx isoform.

**Figure 3.**
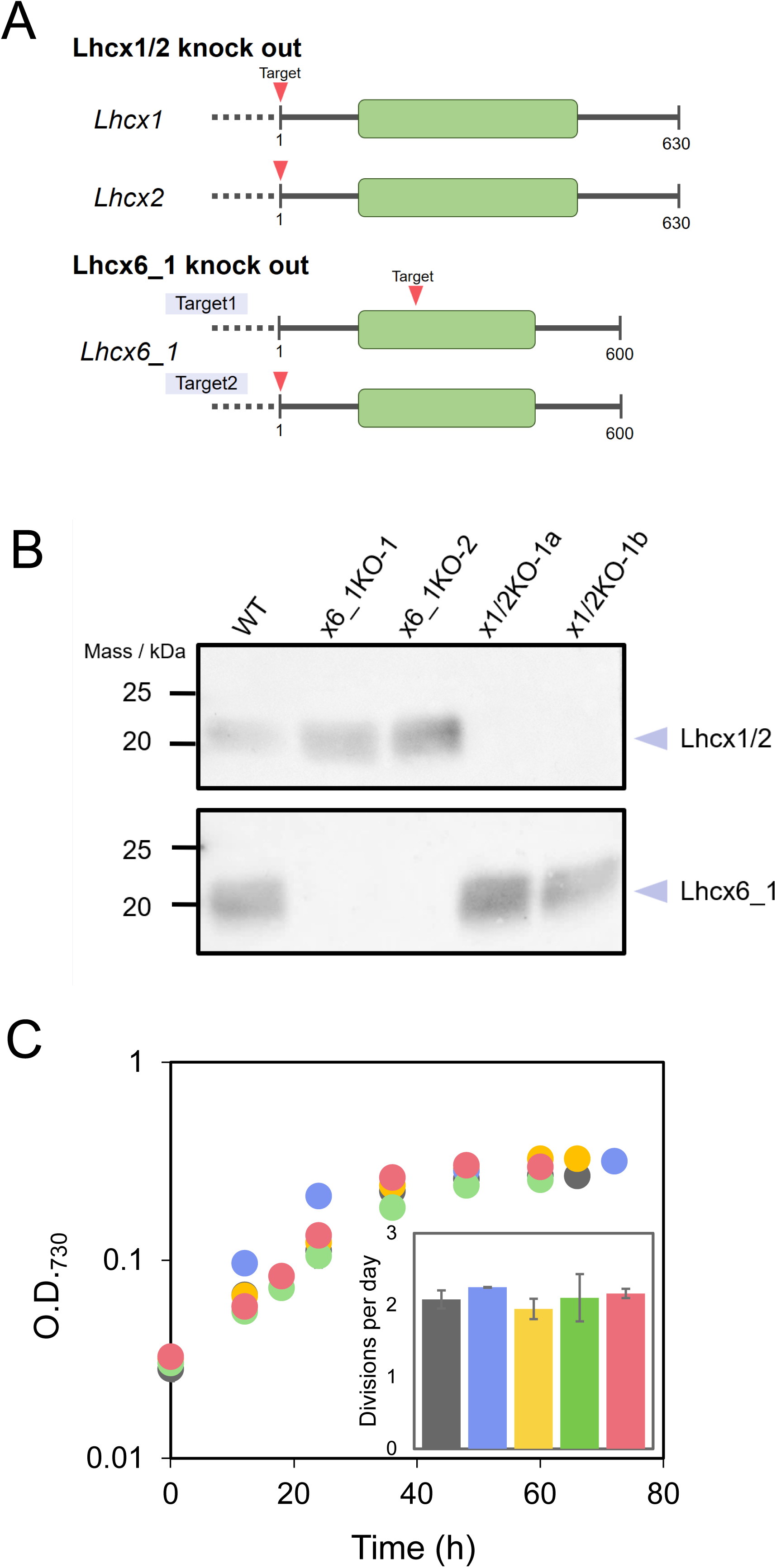
Genome editing of TpLhcx1/2 and TpLhcx6_1 in *Thalassiosira pseudonana*. (A) Target sites for CRISPR/Cas9 nickase in TpLhcx1/2 and TpLhcx6_1. Green bars show chlorophyll binding domains. (B) Immunoblotting of whole crude extracts from the wild type (WT) and genome editing strains (x1/2KOs and x6_1KOs) with antibodies to TpLhcx1/2 and TpLhcx6_1. (C) Growth curves of WT (black), x1/2KOs (blue and yellow), and x6_1KOs (green and red). Divisions per day (inset) was calculated at the logarithmic growth phase. Data are shown as the mean with the standard deviation (*n* = 3, biological replicates).

The cell growth of the established transformants was investigated in comparison with the *T. pseudonana* WT cells, indicating that the deletion of TpLhcx1/2 and TpLhcx6_1 did not affect both the specific growth rate at the logarithmic growth phase (Fig. 3C). The Chl *a* content per cells increased by 130∼150% in these transformants compared with WT (Supplemental Fig. S3A). However, there was no difference in the cell absorption spectra and the ratio of Chl *a* to Chl *c* between WT and each transformant (Supplemental Fig. S3B and C). Further, we analyzed the contents of xanthophyll pigments, including diadinoxanthin, diatoxanthin, violaxanthin, antheraxanthin, and zeaxanthin, based on the Chl *a* content, showing no difference between WT and the transformants (Supplemental Fig. S3D).

### Impact of TpLhcx1/2 and TpLhcx6_1 on photoprotection of PSII

It has been reported that Lhcx isoforms are critical components for qE in the diatom *P. tricornutum*, and the lack of Lhcsr3 leads to PSII photoinhibition in the green alga *C. reinhardtii* (Peers et al. 2009; Bailleul et al. 2010; Buck et al. 2019; Allorent et al. 2013). To assess the impact of TpLhcx1/2 and TpLhcx6_1 on photoprotection in *T. pseudonana*, the maximum efficiency of PSII photochemistry, termed as Fv/Fm, was measured before and after 2-h HL exposure (500 µmol photons m^−2^ s^−1^). Hereafter, these cells were defined as LL cells and HL cells, respectively. While WT showed Fv/Fm at the equivalent level after the HL treatment, both x1/2KO and x6_1KO transformants showed the significant decrease in the values approximately by 30% (Fig. 4A). Photosynthetic net O2 evolution rate was measured before and after the HL exposure, showing the slightly lower maximum photosynthetic activity in x1/2KO and x6_1KO transformants than in the WT in the case of HL cells but not in LL cells (Fig. 4B and C). These results suggested that both Lhcx isoforms are critical to photoprotection of PSII for maintaining photosynthetic activity in HL conditions in *T. pseudonana*.

**Figure 4.**
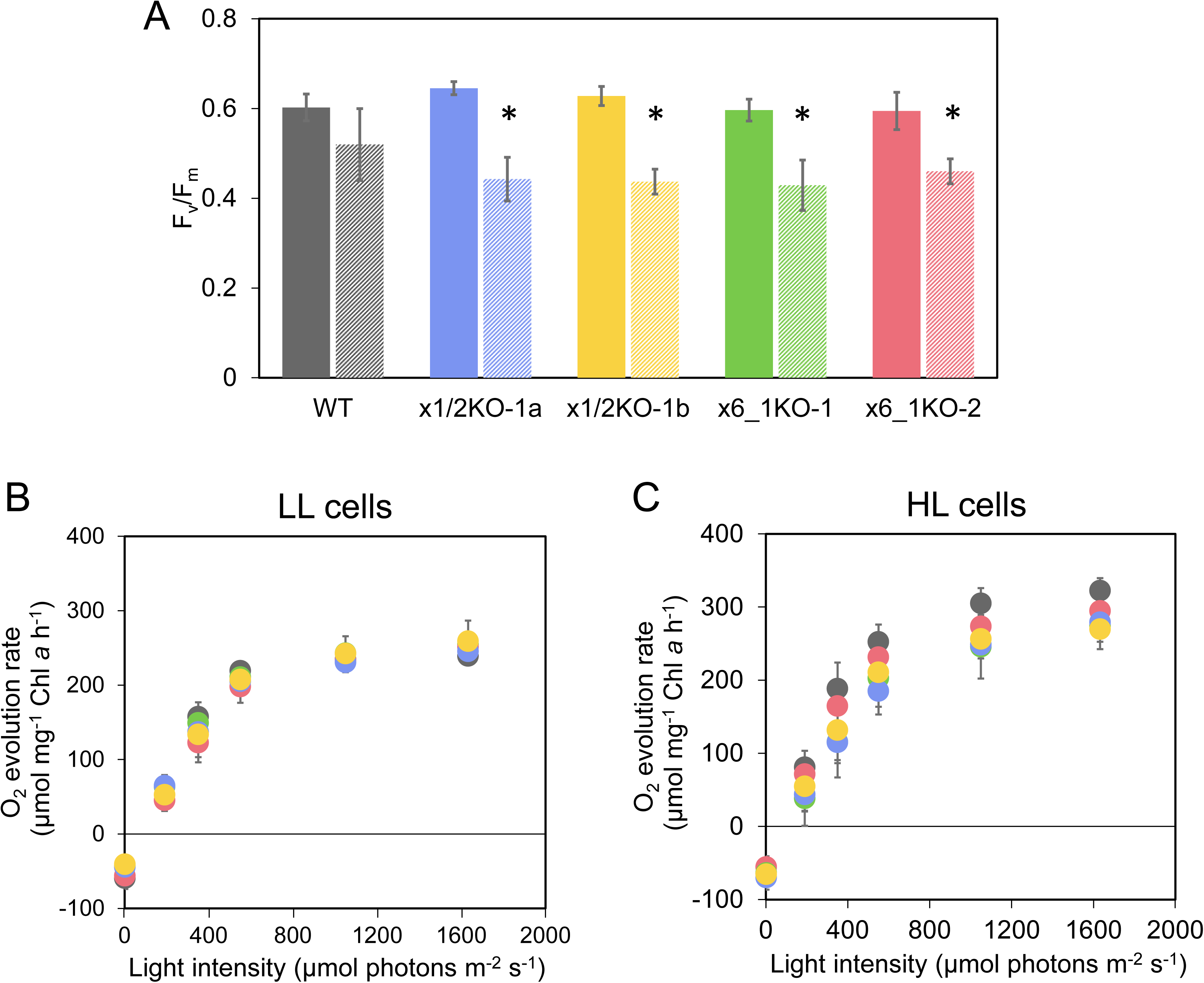
Effects of high light exposure on PSII photochemistry in *Thalassiosira pseudonana* wild type (WT) and genome editing strains (x1/2KOs and x6_1KOs, respectively). Measurements were conducted before and after 2-h exposure to high light (500 µmol photons m^−2^ s^−1^) with 10-min dark interval (LL cells and HL cells, respectively). (A) Maximum quantum yield of PSII photochemistry (Fv/Fm) in WT (black), x1/2KOs (blue and yellow), and x6_1KOs (green and red). Solid and hatched bars show LL and HL cells, respectively. (B, C) Photosynthetic O2 evolution rates at various light intensities in LL cells (B) and HL cells (C) of WT, x1/2KOs, and x6_1KOs. Data are shown as the mean with the standard deviation (*n* = 3, biological replicates). Asterisks indicate the statistically significant differences (*p*<0.05) before and after the high light exposure as per Student’s *t* test.

### Effects of TpLhcx1/2 and TpLhcx6_1 on Chl fluorescence parameters

The *T. pseudonana* WT and Lhcx transformants were further analyzed by pulse-amplitude modulation Chl fluorometry to investigate the molecular mechanisms of PSII photoprotection by TpLhcx1/2 and TpLhcx6_1. In the diatom *P. tricornutum*, the transformants defective of Lhcx isoforms exhibited the impaired capacity of qE, as reflected in the decrease in NPQ during strong irradiance (Buck et al. 2019). Here, we applied a strong actinic light (1180 µmol photons m^−2^ s^−1^) to the *T. pseudonana* WT and transformant cells in the Chl fluorescence measurement for the cells without and with the 2-h HL exposure. In the LL cells of WT, NPQ started to be gradually induced within 4 min after illuminating with the actinic light and finally reach approximately 0.5 in 15 min (Fig. 5A). HL cells of WT showed the similar trend, but NPQ was induced much faster within a few minutes to reach the maximum level (about 0.4; Fig. 5B) that was lower than that of the LL cells. The LL cells of x1/2KO transformants showed the slower NPQ induction than WT irrespective of growth light strength (Fig. 5A and B), and interestingly the HL cells showed much lower maximum NPQ relative to the LL transformants (Fig. 5A and B). These results suggest that TpLhcx1/2 mainly function in qE in *T. pseudonana* resembling the case of *P. tricornutum*. Unexpectedly, x6_1KO transformants did not show the decrease in NPQ, but rather the values were higher than those in WT in both LL and HL cells (Fig. 5A and B). The enhancement of NPQ in x6_1KO transformants was observed especially just after the onset of the illumination in HL cells (Fig. 5B).

**Figure 5.**
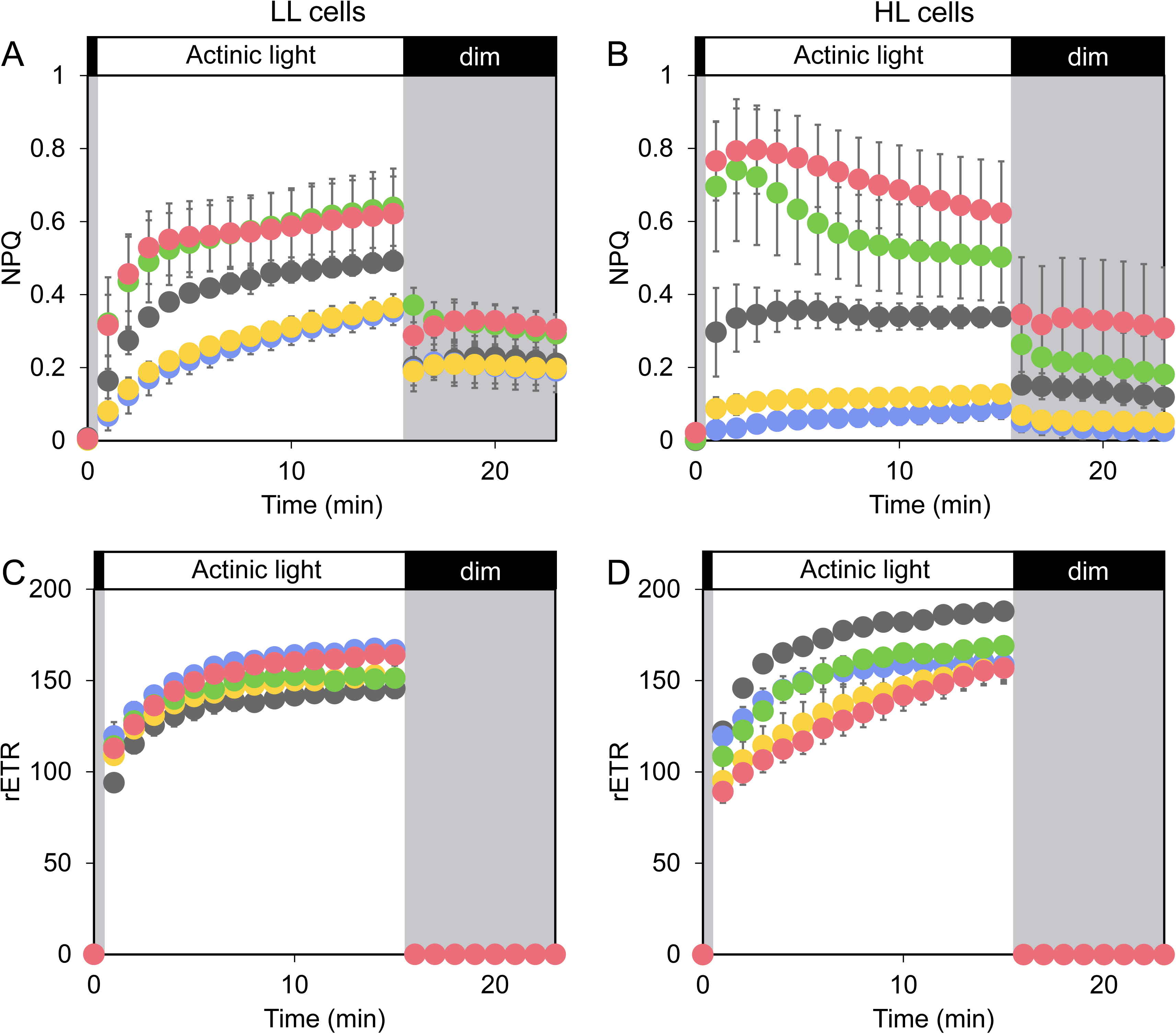
Non-photochemical quenching (NPQ; A and B) and relative electron transport rate at PSII (rETR; C and D) in *Thalassiosira pseudonana* wild type (WT) and genome editing strains (x1/2KOs and x6_1KOs, respectively) before and after 2-h high light exposure (LL cells and HL cells, respectively). After actinic light illumination (1180 µmol photons m^−2^ s^−1^, white bars), the cells were exposed to a dim light (3 µmol photons m^−2^ s^−1^, grey bars) WT, black; x1/2KOs, blue and yellow; x6_1KOs, green and red. Data are shown as the mean with the standard deviation (*n* = 3, biological replicates).

Simultaneously with NPQ, the other Chl fluorescence parameters were measured in each cell. Relative electron transport rate (rETR) was calculated as the product of effective quantum yield of PSII and the light intensity, which is an indicator to photosynthetic linear electron flow in diatoms (Shimakawa et al. 2017). While the WT and Lhcx transformants showed rETR at similar levels in LL cells, both x1/2KO and x6_1KO transformants presented the lower values in HL cells (Fig. 5C and D), consistent with the results of net O2 evolution rate (Fig. 4B and C).

### Effects of TpLhcx6_1 on time-resolved Chl fluorescence

Whereas pulse-amplitude modulation Chl fluorometry uncovered that TpLhcx1/2 play a critical role in induction of qE in both LL and HL cells, the physiological function of TpLhcx6_1 was likely to suppress qE and the mechanism underlying this is enigmatic. Therefore, we measured time-resolved Chl fluorescence at 77 K for the WT cells and x6_1KO transformants. For the measurements, the LL-grown diatom cells were immediately frozen in liquid nitrogen before and after the illumination with an actinic light (1180 µmol photons m^−2^ s^−1^) for 10 min as shown in Fig. 5A. These cell samples were defined as “dark state” and “light state”, respectively. The Chl fluorescence rise and decay curves at 77 K were analyzed, demonstrating that the averaged fluorescence lifetimes were longer in the x6_1KO transformants at the wide range of emission wavelengths, interestingly in both dark and light states (Fig. 6, upper windows). It was found from fluorescence decay-associated spectra that the longer fluorescence lifetimes are mainly derived from the components with 1.4-ns and 4.0-ns time constants around 685−690 nm (A4 and A5, respectively, Fig. 6). These data suggested that TpLhcx6_1 functions in dissipating excess light energy.

**Figure 6.**
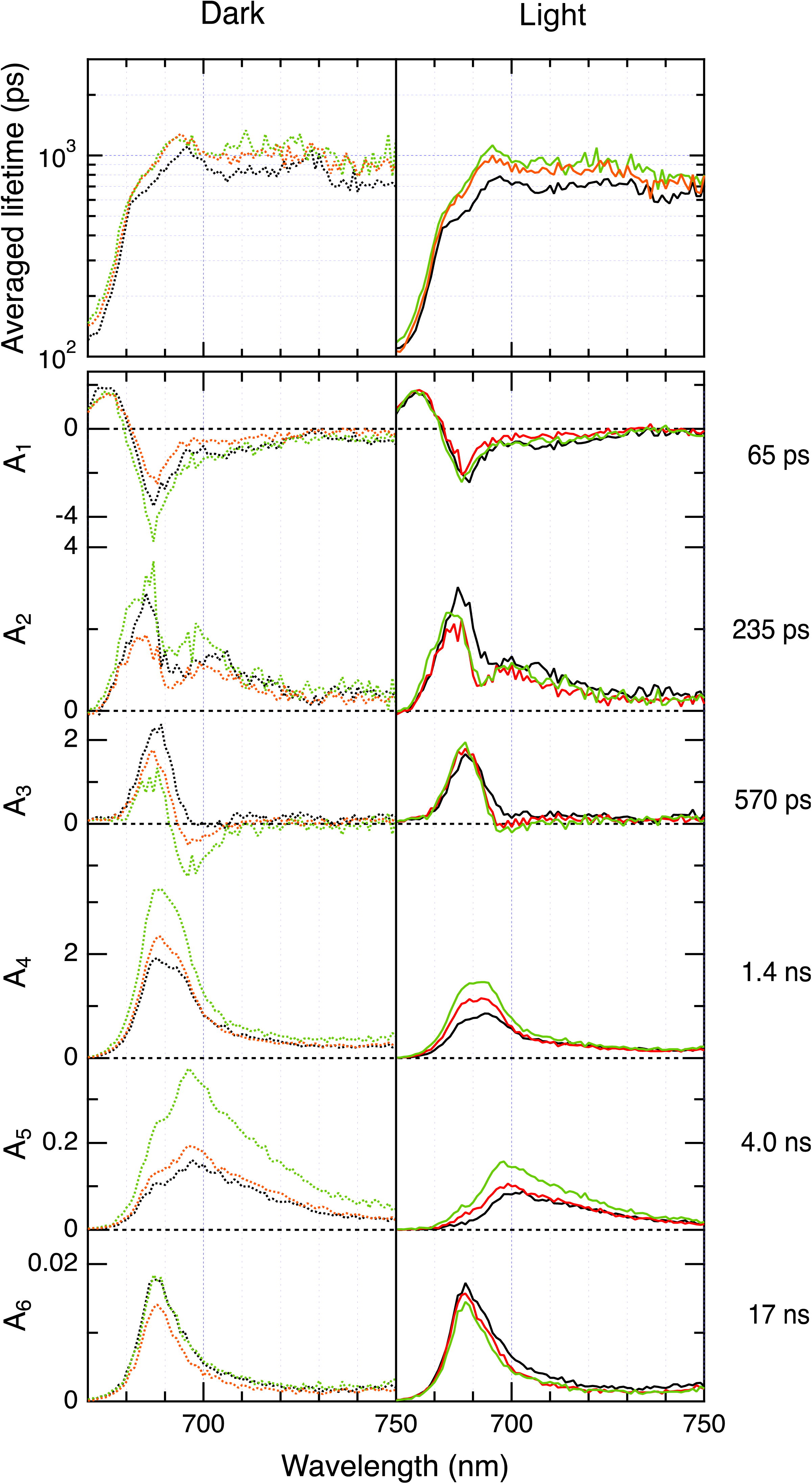
Chlorophyll-fluorescence lifetime and decay-associated spectra at 77 K in *Thalassiosira pseudonana* wild type (WT) and TpLhcx6_1 knock-out transformants (x6_1KOs). Averaged fluorescence lifetime spectra (upper windows) and fluorescence decay-associated spectra (A1–A6, lower windows) are depicted for WT (black) and x6_1KOs (green and red). Cells were immediately frozen at the dark state (dotted line) and 10 min after the onset of actinic light (1180 µmol photons m^−2^ s^−1^; straight line) as shown in Figure 5. The fluorescence decay-associated spectra were normalized by the sum of amplitudes of each sample.

## Discussion

Ocean surface and coastal intertidal zone are easily exposed to a fluctuating light, in which marine diatoms might need to develop a variety of photoprotective mechanisms. The present study uncovered the physiological functions of two types of Lhcx isoforms in the marine diatom *T. pseudonana*. Our data suggested that TpLhcx1/2 are the main component for qE like Lhcx1−3 in *P. tricornutum* (Bailleul et al. 2010), but in contrast, TpLhcx6_1 functions in photoprotection of PSII in the different manner from qE, which should be the photoprotective mechanism uniquely conserved in Thalassiosirales.

The predominant molecular mechanism of qE in *T. pseudonana* is likely to be similar to that already reported in *P. tricornutum*. The Lhcsr or Lhcx isoforms are widely conserved in eukaryotic algae, including green algae, haptophytes, diatoms, and dinoflagellates. TpLhcx1/2 proteins were highly accumulated in HL conditions (Fig. 2A and B)(Calvaruso et al. 2022), which is more similar to the major Lhcx proteins in *C. meneghiniana* (Gundermann et al. 2019) than those in *P. tricornutum* accumulating constitutively (Taddei et al. 2016). The qE capacity was significantly lower in x1/2KO transformants than in WT even after the cells were exposed to high irradiance for 2 h (Fig. 5A and B), indicating that TpLhcx1/2 stayed as a main component for qE even in HL-acclimated cells although the expression of the other TpLhcx genes might be enhanced (Zhu and Green 2010). Nevertheless, it is still not fully understood how TpLhcx1/2 function for qE in the co-operation with xanthophylls. Although it even does not seem to be clear in diatoms how much qE depends on ΔpH of the stroma thylakoid membranes, qE is possibly affected by thylakoid HCO3^−^ transporter and the luminal HCO3^−^ dehydration in the process of CO2-concentrating mechanism (Mukherjee et al. 2019; Nigishi et al. 2024).

Our results suggested that TpLhcx6_1 functions for photoprotection in a different way from the conventional model of qE mechanism. Lhcx6_1 isoforms were phylogenetically grouped into a different clade from Lhcx or Lhcsr, which is conserved uniquely in the Thalassiosirales (Fig. 1). Nevertheless, the gene products have been often detected as the PSII-, PSI-, and FCP-binding proteins in mass spectrometry (Grouneva et al. 2011; Calvaruso et al. 2020), implying that Lhcx6_1 proteins are abundant in these species. Based on the recently-determined structure of PSII-FCP supercomplex, TpLhcx6_1 harbors 11 Chls and 3 carotenoids, which is bound to PSII. TpLhcx6_1 is positioned between PSII and the FCP dimer composed of two TpLhcf7, forming the excitation-energy transfer pathways from the FCP dimer to PSII *via* Lhcx6_1 at short distances. Further, Feng and co-workers speculated that one carotenoid (designated as diatoxanthin 303) in TpLhcx6_1 is an efficient energy quenching site because it was located in close proximity with the Chl clusters but deviated from the efficient excitation-energy transfer pathways from the FCP dimer to PSII (Feng et al. 2023). The x6_1KO transformants showed PSII photodamage as well as x1/2KO ones after the exposure to high irradiance (Fig. 4), indicating that TpLhcx6_1 functions for photoprotection of PSII. However, the deletion of it did not decrease the qE capacity but rather increased, while x1/2KO transformants showed the significantly lower qE capacity, compared with WT (Fig. 5A and B). Importantly, the fluorescence lifetime was prolonged in x6_1KO transformants at low-temperature time-resolved fluorescence in both dark and light states of the cell samples (Fig. 6A and B). Overall, TpLhcx6_1-mediated non-photochemical energy dissipation was constitutively active even in the dark-acclimated cells. In pulse-amplitude modulation fluorometry, qE capacity was evaluated as NPQ that is activated by high irradiance (Baker 2008), which might be the reason why TpLhcx6_1-mediated non-photochemical energy dissipation was not detected as NPQ in that measurement (Fig. 5A and B). Further, x6_1KO transformants showed the higher qE capacity than the WT cells (Fig. 5A and B). Possibly, qE complementarily functioned for photoprotection in the x6_1KO transformants. Based on the preceding biochemical and structural studies, the energy quenching site *via* TpLhcx6_1 should be close to PSII, presumably between CP47 and the FCP dimer (Feng et al. 2023), which is also supported by the results of fluorescence decay associated spectra (Fig. 6). Such a quenching site close to PSII is fitted to the reported site for the energy quenching that has been defined as “qE1a quenching” at a phenomenological level in *C. meneghiniana* (Gundermann et al. 2019; Chukhutsina et al. 2014). Unfortunately, it still remains to assign TpLhcx6_1 as the exact molecular factor for the qE1 quenching at the present time. Our results suggested that the energy quenching *via* TpLhcx6_1 does not seem to be a light-inducible regulatory mechanism like qE but more like a constitutively active system to prevent an “overflow” of excitation energy at PSII-FCP supercomplex.

## Materials and Methods

### Cultures

The marine diatom *T. pseudonana* (Hustedt) Hasle et Heimdal (CCMP 1335) was grown axenically and photoautotrophically cultured in artificial seawater medium with the addition of half-strength Guillard’s ‘F’ solution (Guillard 1975; Guillard and Ryther 1962) supplemented with 10 nM sodium selenite under continuous light (20°C, 50 μmol photons m^−2^ s^−1^, fluorescent lamp). The cultures were aerated with ambient air or 1% CO2. Only sodium chloride concentration of the medium was modified to two thirds in this study.

Diatom growth was evaluated as the optical density at 730 nm. Divisions per day was calculated by the previously reported method (Wood et al. 2005).

### Bioinformatics

The similarity between Lhcx homologs was inferred by the Maximum Likelihood method and Le_Gascuel_2008 model (Le and Gascuel 2008). Initial tree(s) for the heuristic search were obtained automatically by applying Neighbor-Join and BioNJ algorithms to a matrix of pairwise distances estimated using the JTT model, and then selecting the topology with superior log likelihood value. A discrete Gamma distribution was used to model evolutionary rate differences among sites. All positions containing gaps and missing data in the amino-acid sequence alignment were eliminated. The analyses were conducted in MEGA X (Kumar et al. 2018).

Taxonomic distribution of the TpLhcx1/2 and TpLhcx6_1 protein sequences in the ocean was queried against the Ocean Gene Atlas v2.0 webserver (https://tara-oceans.mio.osupytheas.fr) (Villar et al. 2018).

### Immunoblotting

The cells harvested at the logarithmic growth phase were resuspended in a lysis buffer (50 mM Tris-HCl, 2% SDS, pH 6.8) and incubated for 30 min at room temperature. After the Chl *a* quantification, the lysates were applied to SDS-PAGE and immunoblotting. A rabbit anti-TpLhcx1/2 antiserum (diluted 1:2000) targeted to the 197−210 region of the amino acid sequence of TpLhcx1 and the 196-209 region of that of TpLhcx2, and a rabbit anti-TpLhcx6_1 antiserum (diluted 1:2000) targeted to the 133−151 region of the amino acid sequence of TpLhcx6_1 (Japan Bio Serum, Hiroshima, Japan). After incubated with a goat anti-rabbit IgG conjugated with horseradish peroxidase (diluted 1:10000), immunoreactive signals were detected by an enhanced chemiluminescence reagent (ImmunoStar Zeta, Wako, Osaka, Japan) with a high sensitivity CCD imaging system (Luminograph I, ATTO, Tokyo, Japan). Semi-quantitative comparison of band intensity was performed by densitometry with ImageJ. RbcL was detected as a loading control.

### Genome editing of *TpLhcx1/2* and *TpLhcx6_1*

Pairs of sgRNA sequences were designed to target *Lhcx1/2* and *Lhcx6_1* using ZiFiT Targeter (http://zifit.partners.org/ZiFiT/) (Sander et al. 2010), followed by Cas-Designer (http://www.rgenome.net/cas-designer/), which considers specificity and microhomology-mediated joining (Park et al. 2015). The nucleotides 5ʹ-AAACACTCTTCACAATGTTC-3ʹ and 5ʹ-CTCGCCCTCCTCTCCCTCATC-3ʹ were chosen for *Lhcx1/2*, and 5ʹ-CCAACACACCACCATGAAGC-3ʹ and 5ʹ-ATCATCAGTACTCTTATCGC-3ʹ (target 1), and 5ʹ-AAGGCCAGCCCCGAGGAGCTT-3ʹ and 5ʹ-CGTGAGGTTGAGATCATGCAC-3ʹ (target 2) were chosen for *Lhcx6_1* (Fig. 4A and Supplemental Fig. S2). The dual sgRNA vector was generated and introduced together with Cas9 (D10A) nickase vector into the *T. pseudonana* cells by particle bombardment according to the previous work (Nawaly et al. 2020). The transformant strains were screened on agar medium supplemented with 100 µg/mL nourseothricin (Jena Bioscience, Jena, Germany). Monoclonalization of the transformant strains was achieved after four-times respreading and direct sequencing (Supplemental Fig. S2).

### Pigment analyses

For a routine Chl determination, diatom cells were centrifugally harvested, disrupted by vortexing in 100% (v/v) dimethylformamide, and then diluted with 100% (v/v) methanol to 1 mL. Final concentration of dimethylformamide was 10% (v/v). The suspensions were then centrifuged at 10,000 ×*g* for 2 min, and the Chl *a* and *c* contents of the supernatants were determined spectrophotometrically (Jeffrey and Humphrey 1975).

Xanthophyll pigments were extracted from centrifugally harvested cells by vortexing in 100% (v/v) acetone. Then, the extracts were concentrated *via* vacuum centrifugation, and finally the pellet was resuspended in 0.5 M ammonium acetate/methanol at a ratio of 12/88 (v/v). Aliquots (10 µL each) of extracts were applied to the HPLC system, which is composed of a LC-40B X3 pump and SPD-M40 photodiode array detector (Shimadzu, Kyoto, Japan) with an HPLC column (SunShell C18 2.6 µm; ChromaNik Technologies, Osaka, Japan). Detector wavelength was set to 440 nm. Eluent A consisted of 0.5 M ammonium acetate/methanol at a ratio of 15/85 (v/v), and eluent B was methanol/ethyl acetate at a ratio of 90/10 (v/v). The flow rate was adjusted to 0.6 mL min^−1^, and the gradient was run as follows: 3 min of isocratic flow with 100% eluent A followed by a linear gradient to 100% eluent B within 4 min and 5 min of isocratic flow with 100% eluent B, followed by another 3 min for column re-equilibration. Each pigment was determined following the peaks of phytoplankton pigment standards (DHI, Hørsholm, Denmark). Chl *a* was determined simultaneously with xanthophyll pigments in this system, and the xanthophyll pigment concentrations were calculated based on the Chl *a* content.

### Measurements of O2

The concentration of O2 in the medium containing the diatom cells (5 µg Chl *a* mL^−1^) and 10 mM NaHCO3 was monitored with an O2 electrode (DW2/2; Hansatech Ltd, King’s Lynn, UK) at 20°C. Cells were illuminated with white actinic light at various intensities from a halogen lamp (LM-150, Moritex, Saitama, Japan) in a light source Mega Light 100 (Schott, Mainz, Germany). During the measurement, the reaction mixture was stirred with a magnetic stirrer.

### Pulse-amplitude modulation Chl fluorescence analysis

Pulse-modulated excitation was achieved using an LED lamp with a peak emission of 625 nm in a Multi-Color-PAM (Walz, Effeltrich, Germany). Pulse-modulated fluorescence was detected within the range of wavelength limited by RG 665 long pass and SP 710 short pass filters. Cells were illuminated with white actinic light from the LED array. Fv/Fm was calculated as (Fm – Fo)/Fm, NPQ was calculated as (Fm – Fmʹ)/Fmʹ, and effective quantum yield of PSII was calculated as (Fmʹ – Fʹ)/Fmʹ with: Fm, maximum fluorescence from dark-acclimated cells; Fmʹ, maximum fluorescence from light-acclimated cells; Fʹ, fluorescence emission from light-acclimated cells. Short saturation flashes (10,000 µmol photons m^−2^ s^−1^, 600 ms) were applied to determine Fm and Fmʹ.

### Low-temperature time-resolved Chl fluorescence analysis

Fluorescence kinetics were observed by a time-correlated single photon counting system with time and wavelength intervals of 18.3 ps/channel and 1 nm/channel, respectively. The excitation source was a pulse diode laser emitting at 459 nm (PiL047X; Advanced Laser Diode Systems, Berlin, Germany). The observed fluorescence rise and decay curves, *I*_FL_(*λ, t*), were globally analyzed as,

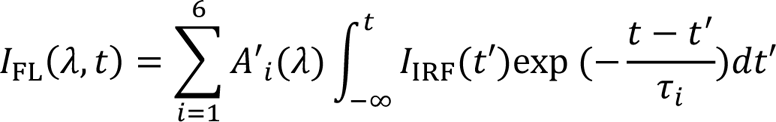

where *I*_IRF_(*t*) and *τ_i_* are the instrumental response function and the *i*th time constant, respectively. The *i*th fluorescence decay-associated spectrum is obtained as *A*′_i_(*λ*), and the normalized fluorescence decay-associated spectrum, *A*′_i_(*λ*), was given using the following equation.

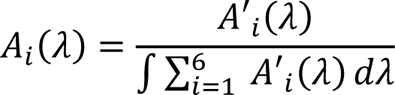

An averaged-lifetime spectrum, 〈*τ*(*λ*)〉, was calculated as,

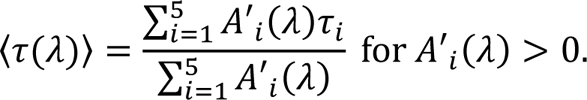

The 6th component is the delayed fluorescence; therefore, this component was excluded in the calculation of 〈*τ*(*λ*)〉.

## Acknowledgements

We wish to thank Mrs. Nobuko Higashiuchi and Mrs. Eri Nakayama (Kwansei-Gakuin University) for their technical assistance, and thank Ms. Miyu Furutani and Ms. Nozomi Sakai (Kobe University) for their help in analyzing fluorescence kinetics.

## Supplementary materials

**Supplemental Table S1.**
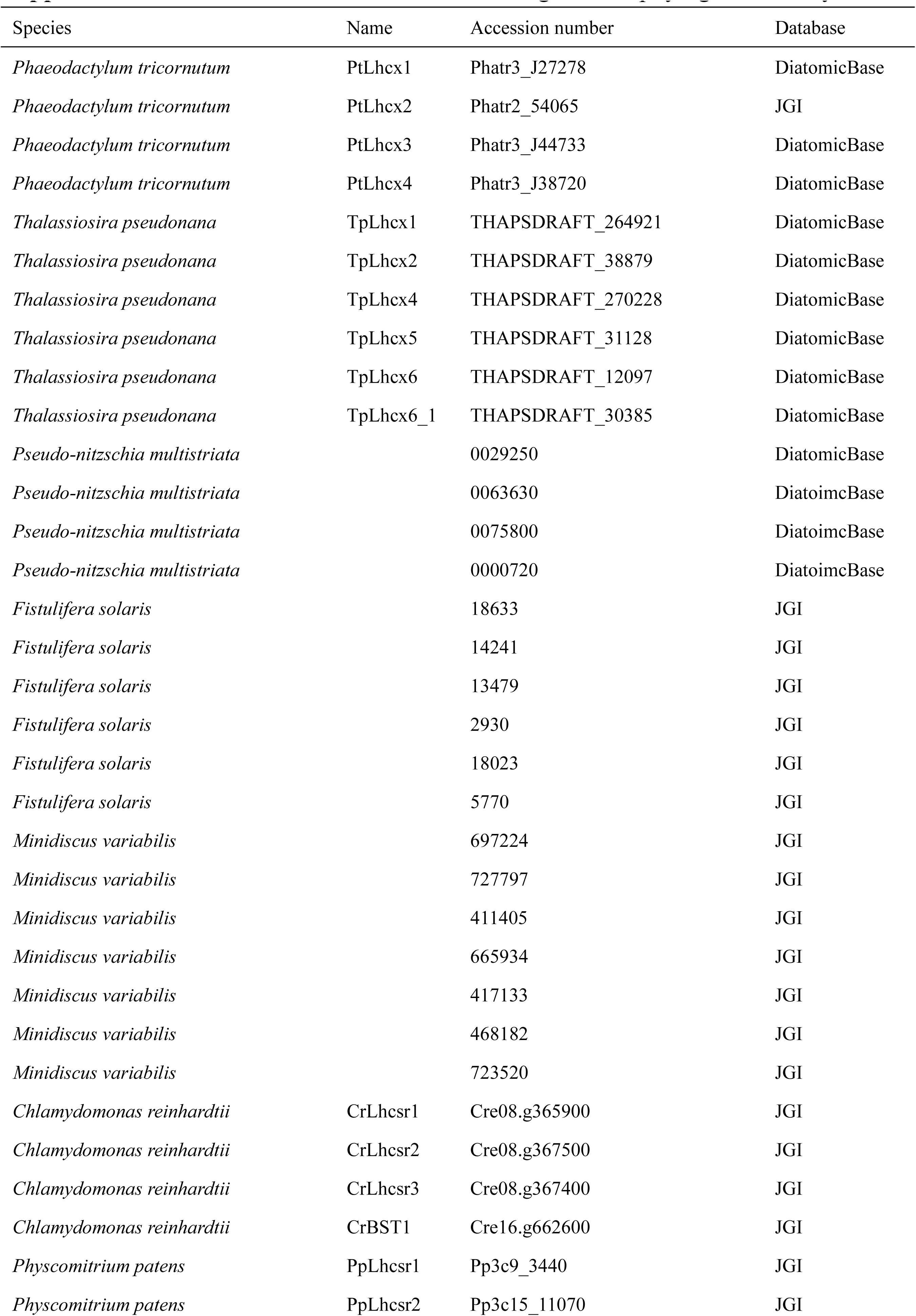
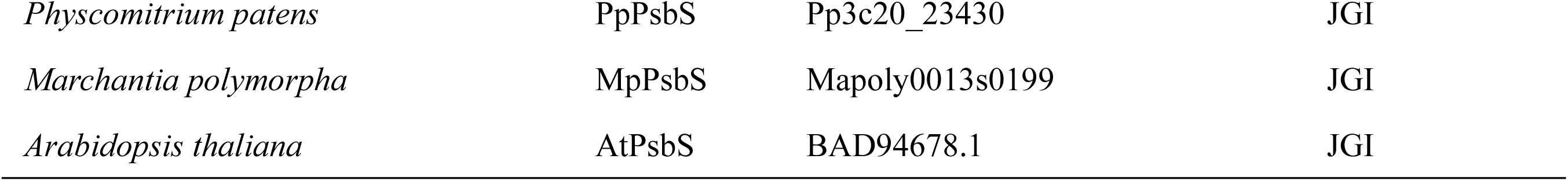
Accession numbers of the genes for phylogenetic analyses.

| Species | Name | Accession number | Database |
| --- | --- | --- | --- |
| *Phaeodactylum tricornutum* | PtLhcx1 | Phatr3_J27278 | DiatomicBase |
| *Phaeodactylum tricornutum* | PtLhcx2 | Phatr2_54065 | JGI |
| *Phaeodactylum tricornutum* | PtLhcx3 | Phatr3_J44733 | DiatomicBase |
| *Phaeodactylum tricornutum* | PtLhcx4 | Phatr3_J38720 | DiatomicBase |
| *Thalassiosira pseudonana* | TpLhcx1 | THAPSDRAFT_264921 | DiatomicBase |
| *Thalassiosira pseudonana* | TpLhcx2 | THAPSDRAFT_38879 | DiatomicBase |
| *Thalassiosira pseudonana* | TpLhcx4 | THAPSDRAFT_270228 | DiatomicBase |
| *Thalassiosira pseudonana* | TpLhcx5 | THAPSDRAFT_31128 | DiatomicBase |
| *Thalassiosira pseudonana* | TpLhcx6 | THAPSDRAFT_12097 | DiatomicBase |
| *Thalassiosira pseudonana* | TpLhcx6_1 | THAPSDRAFT_30385 | DiatomicBase |
| *Pseudo-nitzschia multistriata* |  | 0029250 | DiatomicBase |
| *Pseudo-nitzschia multistriata* |  | 0063630 | DiatomicBase |
| *Pseudo-nitzschia multistriata* |  | 0075800 | DiatomicBase |
| *Pseudo-nitzschia multistriata* |  | 0000720 | DiatomicBase |
| *Fistulifera solaris* |  | 18633 | JGI |
| *Fistulifera solaris* |  | 14241 | JGI |
| *Fistulifera solaris* |  | 13479 | JGI |
| *Fistulifera solaris* |  | 2930 | JGI |
| *Fistulifera solaris* |  | 18023 | JGI |
| *Fistulifera solaris* |  | 5770 | JGI |
| *Minidiscus variabilis* |  | 697224 | JGI |
| *Minidiscus variabilis* |  | 727797 | JGI |
| *Minidiscus variabilis* |  | 411405 | JGI |
| *Minidiscus variabilis* |  | 665934 | JGI |
| *Minidiscus variabilis* |  | 417133 | JGI |
| *Minidiscus variabilis* |  | 468182 | JGI |
| *Minidiscus variabilis* |  | 723520 | JGI |
| *Chlamydomonas reinhardtii* | CrLhcsr1 | Cre08.g365900 | JGI |
| *Chlamydomonas reinhardtii* | CrLhcsr2 | Cre08.g367500 | JGI |
| *Chlamydomonas reinhardtii* | CrLhcsr3 | Cre08.g367400 | JGI |
| *Chlamydomonas reinhardtii* | CrBST1 | Cre16.g662600 | JGI |
| *Physcomitrium patens* | PpLhcsr1 | Pp3c9_3440 | JGI |
| *Physcomitrium patens* | PpLhcsr2 | Pp3c15_11070 | JGI |
| *Physcomitrium patens* | PpPsbS | Pp3c20_23430 | JGI |
| *Marchantia polymorpha* | MpPsbS | Mapoly0013s0199 | JGI |
| *Arabidopsis thaliana* | AtPsbS | BAD94678.1 | JGI |

**Supplemental Figure S1.**
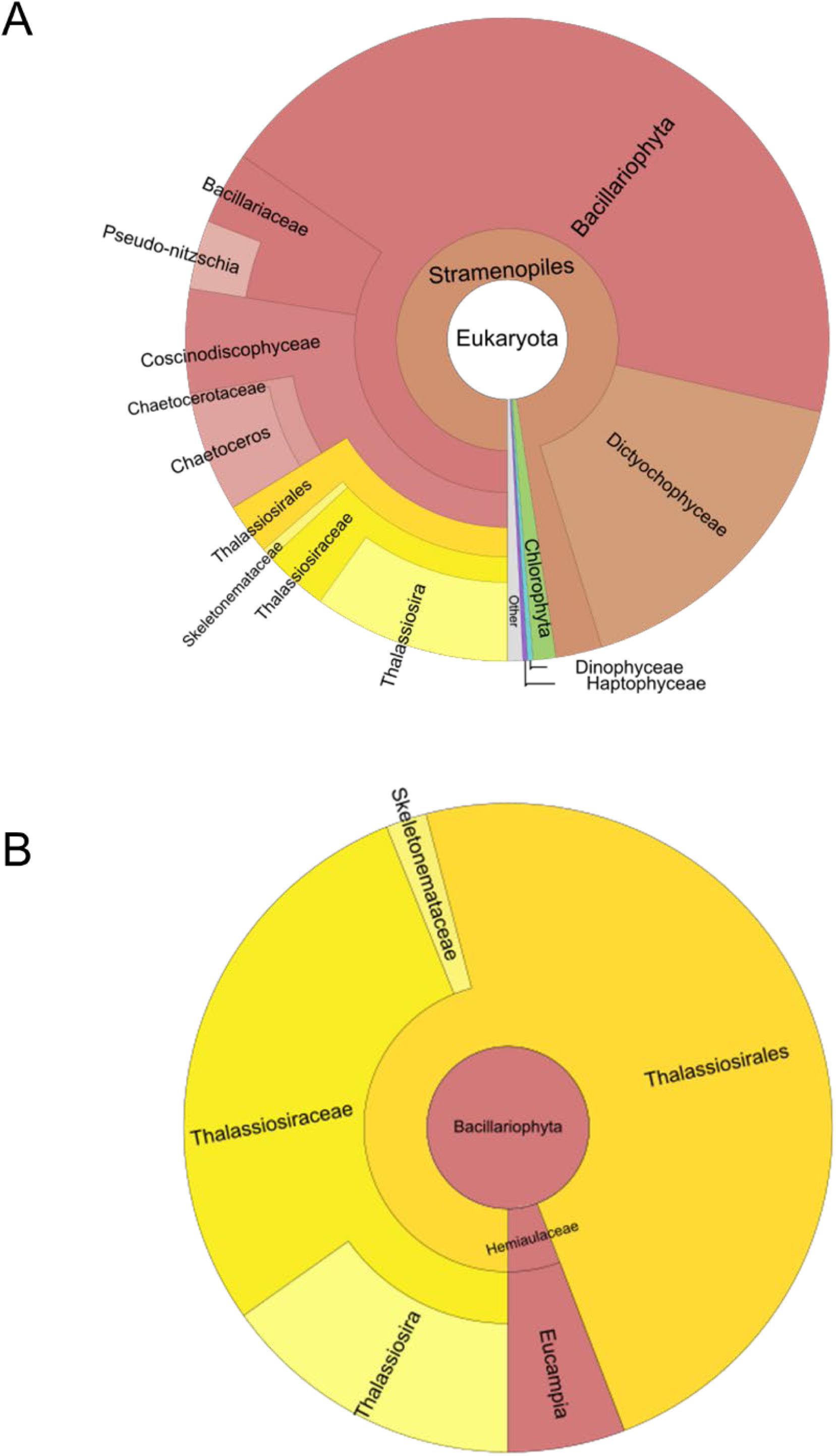
Environmental metagenomics distribution of homologs of TpLhcx1 (A) and TpLhcx6_1 (B) based on the prediction in Ocean Gene Atlas (see the Materials and Methods for the details).

**Supplemental Figure S2.**
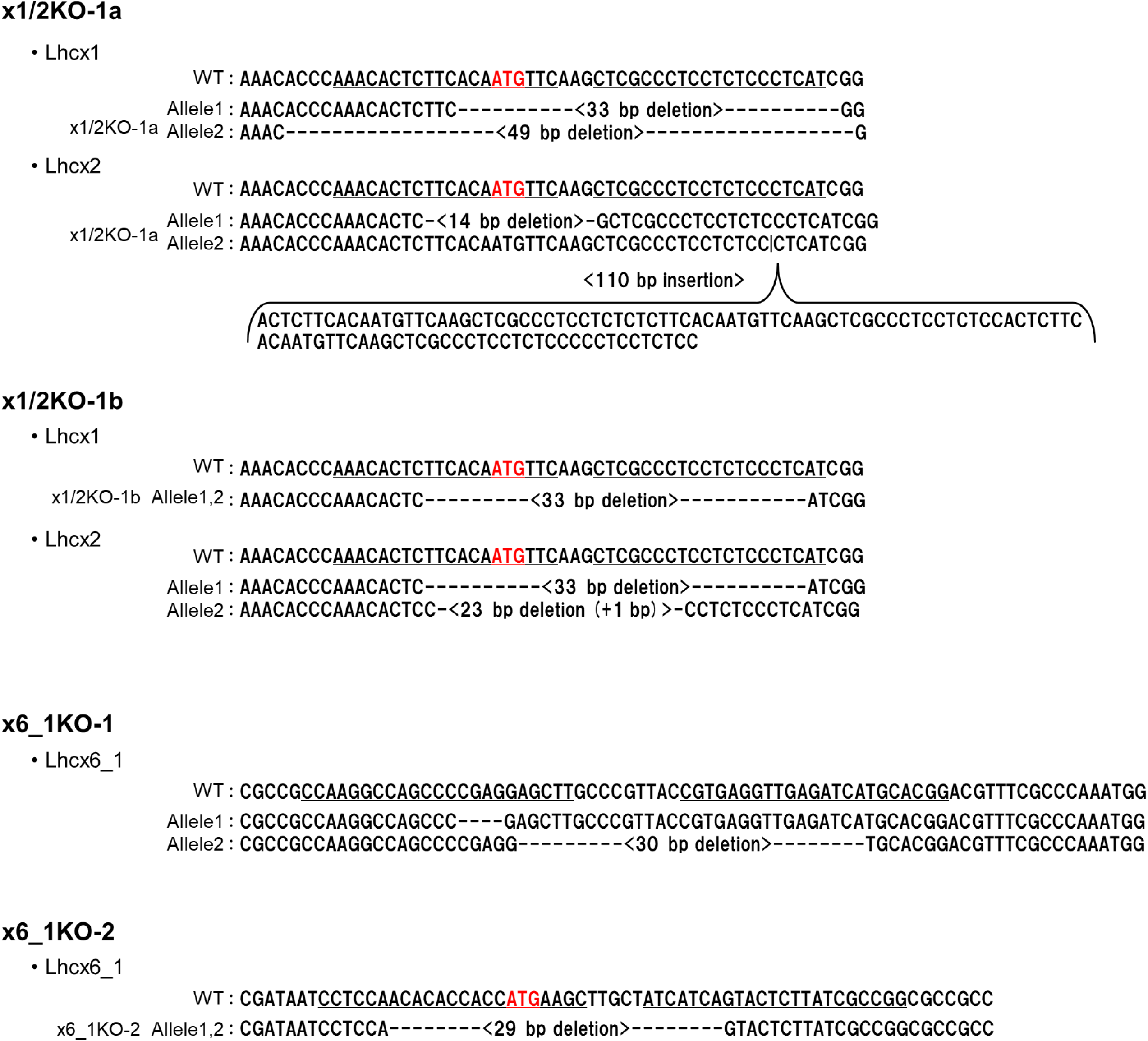
Genome sequences around the genome editing target sites of *TpLhcx1/2* and *TpLhcx6_1* in each knock-out transformant. The initial codons were highlighted in red.

**Supplemental Figure S3.**
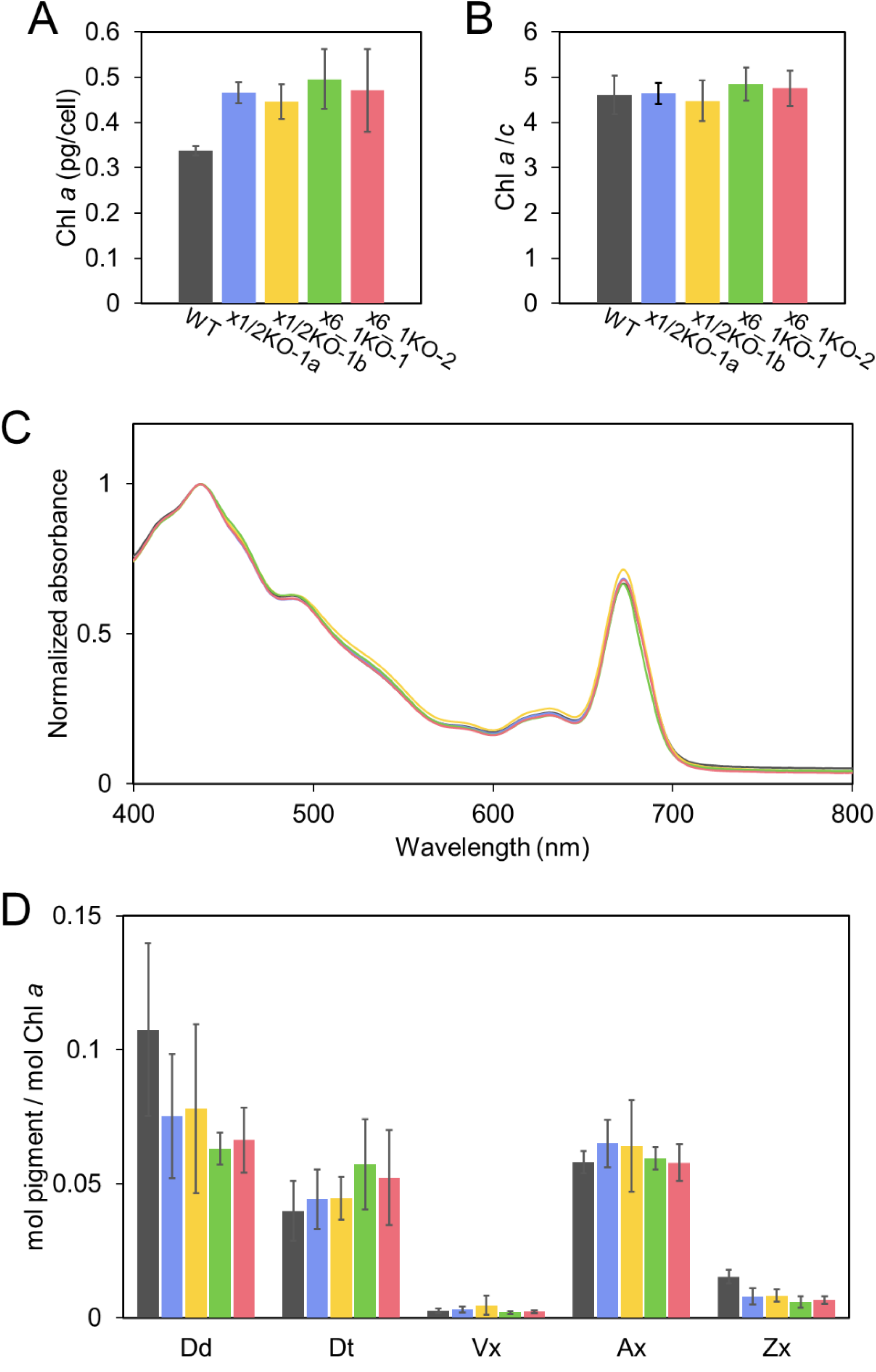
Pigments of *Thalassiosira pseudonana* wild type (WT, black) and genome editing strains (x1/2KOs, blue and yellow; x6_1KOs, green and red). Chlorophyll *a* content (A), chlorophyll *a*/*c* ratio (B), cell absorption spectrum (C), and the contents of xanthophyll pigments (D), including diadinoxanthin (Dd), diatoxanthin (Dt), violaxanthin (Vx), antheraxanthin (Ax), and zeaxanthin (Zx) were determined in the cells grown under low light (50 µmol photons m^−2^ s^−1^). Cell absorption spectra were normalized at the peak of carotenoids. Data are shown as the mean with the standard deviation (*n* = 3, biological replicates).

